# Bile acids accumulate norovirus-like particles and enhance binding to and entry into human enteric epithelial cells

**DOI:** 10.1101/2023.04.28.538707

**Authors:** Elin Palm, Katarina Danskog, Stefan Nord, Miriam Becker, Hugh Tanner, Linda Sandblad, Daniel Öhlund, Annasara Lenman, Niklas Arnberg

**Author notes:** Address correspondence to Niklas Arnberg, or Elin Palm,. These authors contributed equally to this work (Elin Palm, Katarina Danskog). Author order between the two was alphabetical according to first name.

## Abstract

Human norovirus (HuNoV) is a leading cause of acute viral gastroenteritis, but despite high impact on public health and health care, the mechanisms of viral attachment to and entry into target cells are not yet fully understood. It is well known that body fluids such as blood can transmit unrelated viruses, but recent reports also indicate that saliva and bile contribute to transmission of HuNoV. For example, human bile acids increase cell surface ceramide levels in human enteroids, which improves norovirus entry into cells resulting in enhanced replication. Bile acids can also interact directly with the norovirus capsid, but it is not known whether bile or other gastrointestinal body fluids directly affect HuNoV attachment to host cells. In this study, we investigated the effects of patient-derived gastric juice, pancreatic juice, and bile on HuNoV GII.4 virus-like particle (VLP) attachment to and entry into a human duodenal cell line, HuTu-80. We show that while gastric juice and pancreatic juice do not affect viral attachment or entry, bile – in particular hydrophobic bile acids – significantly enhance cellular attachment and subsequent entry of GII.4 VLPs into cells. In addition, we show that hydrophobic bile acids induce accumulation of viral particles in the vicinity of cells. Our results suggest the presence of a new *en masse* infection mechanism, where bile acids aggregate virions, and allow direct and more efficient attachment to and entry into target cells.

**Importance:** Viruses transmitted by the fecal-oral route encounter secreted host factors in gastrointestinal fluids. Some host factors can be exploited by the virus to facilitate infection. Human bile acids indirectly promote norovirus entry into and infection of human enteroids, but the direct effect of bile acids on attachment and uptake, along with the impact of other gastrointestinal fluids, remain unknown. Here, we investigated the direct effects of human body fluids on cellular attachment of norovirus VLPs. We show that human bile and hydrophobic bile acids induce an accumulation of norovirus VLPs, which is associated with significantly enhanced attachment and entry into human duodenal cell lines. These results highlight the differential effects of gastrointestinal body fluids on viral attachment and entry, while providing useful information into the complex HuNoV-host interactions that facilitate infection.

## Introduction

Human norovirus (HuNoV) is one of the most common causes of acute viral gastroenteritis worldwide. It is estimated to cause over 200,000 deaths annually, and constitutes a global economic burden of more than $60 billion (1). Noroviruses are non-enveloped positive-sense RNA viruses belonging to the *Caliciviridae* family. They are classified into 10 different genogroups (GI-GX), which are further divided into 48 genotypes. Viruses belonging to genogroup I, II, IV, VIII, and IX infect humans (2, 3), and out of these genotype GII.4 is currently the most common cause of acute norovirus infection (4–6). The genome is contained within a shell of major capsid protein VP1 and minor capsid protein VP2, where VP2 is located inside the capsid to increase stability. VP1 consists of an S and a P domain (7, 8), where the P domain is further subdivided into a P1 and a receptor interacting P2 domain. The sequence of the P2 domain is the least conserved and contains sites that contribute to cell binding, antigenicity and immune driven evolution (9). Expression of VP1 in the baculoviral-expression system results in self-assembly of virus-like particles (VLPs) that are comparable to the native virus in size and appearance (10).

To date, there are no available vaccines or antivirals, partly due to HuNoV research being hampered by the – until recently (11) - lack of a more user-friendly cell model to cultivate the virus (3, 12–16). Research on HuNoV has thus resorted to using alternatives such as virus-containing stool samples, murine norovirus (MNoV), which readily replicates *in vitro* but does not cause gastrointestinal disease in mice (17), and VLPs that consist of an empty virus capsid with capacity to bind and enter cells (18, 19). Susceptibility to HuNoV infection of certain strains is associated with expression of histo-blood group antigens (HBGAs) on epithelial cells, mediated by the enzyme fucosyltransferase-2 (FUT2) (18, 20–22), where the P2 domain of VP1 binds to fucose on the cell surface. Fucose is an important attachment factor, and so-called secretor-negative individuals lacking a functional fucosyltransferase-encoding *FUT2* gene, are highly resistant to infection with HBGA-dependent norovirus (23, 24). However, some studies have found that secretor-negative individuals can still be infected by HBGA-dependent norovirus genotypes (25–27). In a recent study it was demonstrated that the Shell (S) domain of VP1 binds to galectin 3 and apoptosis-linked gene 2-interacting protein X (ALIX) (28).

It is known that human tissue fluids contain components that affect virus attachment to target cells, including coagulation factors and lactoferrin, which facilitate non-canonical cellular attachment and entry of human adenovirus (29–31), and host trypsin-mediated cleavage of influenza A hemagglutinin, is essential for membrane fusion (32, 33).

For certain HuNoV strains, such as GII.4, human saliva is important for transmission (11), and human bile is essential for successful replication in human intestinal enteroids (5, 34). Bile is a gastrointestinal body fluid mostly consisting of highly hydrophobic bile acids synthesized in the liver from cholesterol (35–37). A recent study demonstrated that bile acid glycochenodeoxycholic acid (GCDCA) increase the production of cell surface ceramide, which results in an increased uptake of HuNoV particles in human intestinal enteroids (34). This suggests that bile acids enhance HuNoV replication via indirect, cellular effects, but structural studies have also identified bile acid binding pockets in a number of HuNoV strains including GII.1, GII.4, GII.10, and GII.19 (38, 39), indicating that bile acids can have multiple roles in the HuNoV replication cycle.

While it is clear that bile contributes to HuNoV infection, the role of other gastrointestinal fluids such as gastric juice and pancreatic juice in HuNoV entry has not been investigated. In this study, we collected patient-derived gastrointestinal fluids and investigated their effect on VLP interactions with duodenal cells. We found that gastric juice and pancreatic juice did not affect the initial steps in the viral replication cycle. Bile acids, and specifically hydrophobic bile acids on the other hand, contributed to attachment and uptake of GII.4 VLPs into HuTu-80 cells. Additionally, we showed that the VLPs form clusters when incubated with bile and highly hydrophobic TLCA. Several enteric viruses, including norovirus, can be released within extracellular vesicles containing multiple viral particles, thereby increasing the chance of effective infection both *in vitro* and *in vivo* (40, 41). The bile acid-dependent accumulation of viral particles demonstrated in this study suggests the presence of another potential mechanism used by HuNoV for attachment to, entry into, and productive infection of duodenal cells.

## Materials and methods

### Cells and antibodies

Human duodenal adenocarcinoma cell line HuTu-80 (ATCC #HTB-40) was cultured in Dulbecco’s Modified Eagle’s Medium (DMEM; Sigma-Aldrich), 20 mM HEPES (Sigma-Aldrich), 10% Fetal Bovine Serum (FBS; Cytiva), 100 units/mL of penicillin + 100 µg/mL of streptomycin (PEST, Gibco). MAB227P Monoclonal antibody to Norovirus (Mouse IgG2a κ, Clone 2002-G5, Maine Biotechnology Services), rabbit anti-Alexa Fluor 488-conjugated polyclonal IgG (Thermo Fisher Scientific), Alexa Fluor 647-conjugated rabbit anti-mouse polyclonal IgG (Thermo Fisher Scientific), Alexa Fluor 568-conjugated goat anti-rabbit IgG (Thermo Fisher Scientific), Hoechst33342 (Thermo Fisher Scientific), Alexa Fluor 647-conjugated Wheat Germ Agglutinin (Thermo Fisher Scientific).

### VLP production

Human norovirus GII.4 VLP, strain NLV/DIJON171/96 (GeneBank accession no. AAL79839.1). VLPs were expressed with the Bac-to-Bac TOPO Expression system (Invitrogen, Catalog nos. A11101, A11100) according to instructions from the manufacturer. Supernatant was collected from infected SF-9 cells and cellular debris were removed by centrifugation at 3000 x g for 30 min at 4°C. The VLPs were concentrated by high speed centrifugation at 100 000 x g for 2 h at 4°C. The obtained pellet was dissolved in 0.2 M Tris-HCl pH 7.4, loaded onto a discontinuous sucrose gradient 20-60% and centrifuged at 175 000 x g for 16 h at 4°C. From the sucrose gradient, the band corresponding to intact VLPs were extracted and stored at −80°C until further use. For use in experiments, the buffer was exchanged to 25 mM HEPES, pH 8.2 by use of an Amicon Ultra Centrifugal filter, 100 kDa MWCO, and VLPs were stored at +4°C.

### Human body fluid samples and bile acids

Gastric juice, pancreatic juice and bile was collected from adults undergoing hepatobiliary surgery at the surgical department at Umeå University Hospital. The study was conducted according to declaration of the Helsinki and was approved by the Swedish Ethical Review Authority (2019–00399 and 2019-03805). All subjects taking part in the study provided written informed consent. Samples were aliquoted and stored at −80°C until use. Bile acids were purchased from Sigma-Aldrich. GUDCA, GCA, GDCA, GCDCA, TCDCA, TCA, TDCA were dissolved in ultrapure water, whereas CA, CDCA, DCA, UDCA, LCA, TLCA were dissolved in dimethyl sulfoxide (DMSO), and cholesterol (Sigma-Aldrich) in ethanol, all to a concentration of 50 mM.

### Binding assay

HuNoV VLP binding was evaluated using flow cytometry. HuTu-80 cells were detached using PBS supplemented with 0.05% EDTA, counted and reactivated in growth medium for 1 h at 37°C on a tipping table. Cells were plated in 96 well plates (2 x 10⁵ cells/well) and washed with PBS. HuNoV GII.4 VLPs were then preincubated with patient-derived body fluids (bile, pancreatic juice or gastric juice) diluted in PBS to a concentration of 12.5%, or with pure bile acids/cholesterol (diluted in PBS to 4 mM), for 30 min at 37°C on a rotary shaker. Before addition to cells, the mixtures were further diluted in PBS to a final concentration of 0.78% or 250 µM, respectively, and a VLP concentration of 6 µg/mL. The VLP mixtures were added to cells and incubated for 1 h at 4°C on a rotary shaker. Unbound VLPs were washed away with PBS. For staining, anti-norovirus antibody diluted 1:750 in FACS buffer (PBS with 2% FBS) was added and incubated with the cells for 30 min at 4°C on a rotary shaker. Cells were washed once with FACS buffer and secondary antibody rabbit anti-Mouse Alexa Fluor 647 (Thermo Fisher Scientific) diluted 1:1000 in FACS buffer was added and incubated with the cells for 30 min at 4°C on a rotary shaker. Cells were washed once and analyzed by flow cytometry with a BD Accuri C6 instrument (BD Biosciences).

For bile acid pretreatment of cells before VLP binding: cells were incubated with bile acids for 30 min at 37°C. Supernatant containing excess bile acids was removed before VLP mixtures were added for 1 h at 4°C on a rotary shaker followed by staining as described above.

### Norovirus VLP labelling

Norovirus VLPs were labelled with an Alexa Fluor 488 NHS Ester (Thermo Fisher Scientific) dissolved in DMSO at a concentration of 10 µg/mL. Dye was added to purified Norovirus VLPs (in 25 mM HEPES buffer pH 8.2) at a 10x molar excess while vortexing, and the reaction was left for one hour at room temperature under constant rotation. Afterwards, excess dye was removed by buffer exchange to PBS using filter centrifugation (Amicon, Millipore).

### Uptake assay

HuTu-80 cells were seeded in 8 well glass bottom chamber slides (Nunc, Lab Tek II) at a density of 150 000 cells/well. The following day Alexa Fluor 488-labelled VLPs (6 µg/well) were pre-incubated with TLCA, GCDCA or human bile (sample B5) at a concentration of 4 mM (bile acids) or 12.5% (bile), at 37°C for 30 minutes and subsequently diluted to 250 µM (0.78 for bile samples) in PBS. VLPs were allowed to bind to the cell surface for one hour at 4°C. Unbound particles were washed away and the remaining were allowed to internalize for 2 hours at 37°C. Binding control samples were kept on ice. Samples were incubated with blocking buffer (PBS supplemented with 2% BSA) for 15 min and stained for non-internalized virus particles with an anti-Alexa Fluor 488 antibody 1:500 in blocking buffer for 1.5 hours and a secondary Alexa Fluor 568 1:1000 in blocking buffer for 1 hour. Cell nuclei were stained using Hoechst33342 1:5000 and Alexa Fluor 647-conjugated Wheat Germ Agglutinin 1:500 diluted in blocking buffer was used for plasma membrane visualization. Samples were imaged with a Leica SP8 Confocal at a magnification of 63x and z-stack images were acquired, image settings were consistent for all samples. Maximum intensity projections were created using the ImageJ distribution Fiji (42). For quantification of virus entry, maximum intensity projections were analyzed using CellProfiler v4.1.3 (43). Virus particles were identified as objects by the Alexa Fluor 488 signal (all virus), and the images were masked for Alexa Fluor 488-based objects, before identification of objects based on the Alexa Fluor 568 signal (virus outside). Object counts were summarized using pivot tables and virus uptake was measured as a relative comparison between Alexa Fluor 568 and Alexa Fluor 488 signal in each sample, and was shown as fold increase in uptake samples compared to the average background uptake of the corresponding binding control samples. Statistical analysis of the fold increase was performed using Welch’s *t* test. Due to a very low amount of total signal in samples with only VLP, this condition was omitted for uptake quantification. Number of cells were determined by manual counting from maximum intensity projections. More than 70 cells and at least 3 fields of view per condition were imaged and analyzed.

### Electron microscopy

Bile acids TLCA, GCDCA or human bile were mixed with 15 µg HuNoV VLPs in 20 mM HEPES buffer pH 8.2 to a final concentration of 250 µM or 4 mM, followed by 30 minutes incubation at 37°C. For negative staining electron microscopy (EM) imaging, copper grids were coated with thin layers of formvar and carbon (sputtered with Leica EM ACE 200), and glow discharged (PELCO easiGlow). 3.5 µL VLP samples were applied to each grid, blotted to filter paper, washed with two drops of 50 µL H_2_0, stained with a 50 µL drop of 1.5% Uranyl Acetate on parafilm. Stained VLPs were image with Talos L120 transmission electron microscope (TEM, Thermo Fisher Scientific, former FEI) with 47000x magnification. Micrographs were acquired with Ceta 4k x 4k CMOS detector using TEM Image & Analysis software (Thermo Fisher Scientific, former FEI).

### Statistical analysis

All experiments were performed three times in duplicates or triplicates. The results are shown as mean ± SD. Two way-analysis of variance (ANOVA), Student’s *t* test or Welch’s *t* test were performed using GraphPad Prism version 9.3 for Windows (GraphPad Software, San Diego, California USA). A *P*-value of <0.05 was considered statistically significant.

## Results

### HuNoV VLPs can bind to human duodenal cells in presence of human bile or hydrophobic bile acids

It has previously been demonstrated that bile acids increase cell surface ceramide expression resulting in enhanced uptake of HuNoV into cells (34). Since bile acids interact directly with the norovirus capsid, we investigated whether bile, but also other human gastrointestinal fluids affect direct cell attachment of HuNoV GII.4 VLPs. For this purpose, we collected samples of human gastric juice, pancreatic juice, and bile from patients undergoing gastrointestinal surgery. VLPs (6 µg/well) were incubated with gastrointestinal fluids at a concentration of 12,5% (diluted in PBS) and their subsequent attachment to a human duodenal adenocarcinoma cell line (HuTu-80) was quantified by flow cytometry. VLP binding was unaffected by gastric juice (Fig. 1A) and pancreatic juice (Fig. 1B) samples but was largely enhanced by some – but not all – bile samples (Fig. 1C). Thus, we next investigated whether bile acids were able to directly affect binding of VLPs to cells. We pre-incubated VLPs with individual bile acids and quantified binding to HuTu-80 cells. Several bile acids increased binding, among them GCDCA, which has previously been associated with increased HuNoV replication in enteroids (34) (Fig. 2A). We also observed a very strong (30-fold) increase in attachment by highly hydrophobic taurine-conjugated TLCA. These results are in concordance with previous findings, which showed a general selectivity of HuNoV towards hydrophobic bile acids in relation to cell uptake and replication, even though many bile acids are structurally very similar (Fig. 2B) (34).

**Figure 1.**
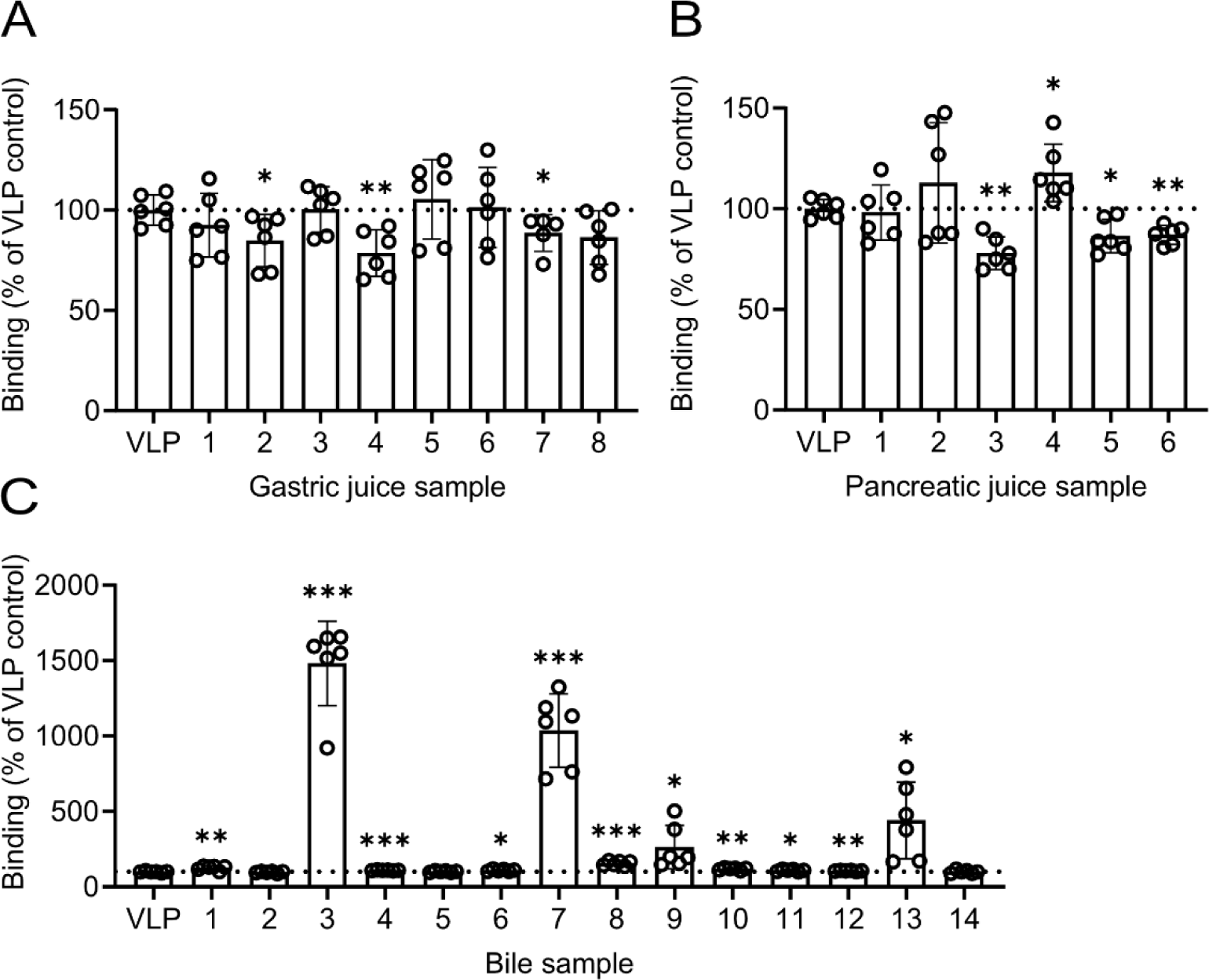
Flow cytometry analysis of HuNoV GII.4 VLP binding to HuTu-80 cells in the presence of human gastrointestinal fluids. VLP binding to HuTu-80 cells in the presence of gastric juice (A), pancreatic juice (B) or bile (C) samples from patients. Binding was measured as median fluorescence and presented as percent of control (VLP binding in absence of human gastrointestinal fluids). Data was generated from three independent experiments, represented as means ± SD Statistical significance was determined with Student’s *t* test; *, P < 0.05; **, P < 0.002; *** P, < 0.001.

**Figure 2.**
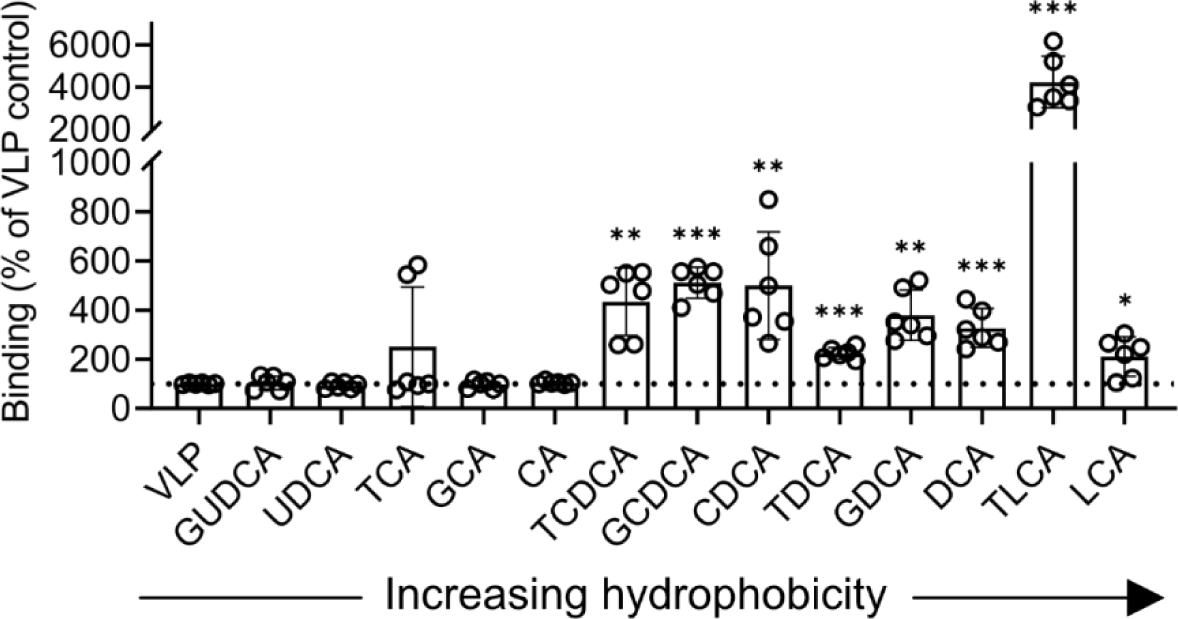
Flow cytometry analysis of HuNoV GII.4 VLP binding to HuTu-80 cells in the presence of bile acids. VLP binding to HuTu-80 cells in the presence of bile acids, arranged in order of increasing hydrophobicity (from left to right). Binding was measured as median fluorescence and presented as percent of control (VLP binding in absence of bile acids). Data are from three independent experiments, represented as means ± SD. Statistical significance was determined with Student’s *t* test; *, P < 0.05; **, P < 0.002; *** P, < 0.001.

### Bile acids exert a direct effect on HuNoV VLP binding to cells

We pre-incubated VLPs or cells with bile acids prior to incubation of VLPs and cells to further investigate the nature of bile acid-mediated binding. When cells were pre-incubated with bile acids prior to addition of VLPs only TLCA significantly increased VLP binding (Fig. 3A “Cell pre-incubation”). However, VLP binding was dramatically increased when the particles were preincubated with TLCA prior to incubation with cells (Fig. 3A “VLP pre-incubation”). This indicates that the increased VLP binding seen in Fig. 2A was not due to bile acid-induced cellular changes but rather required a direct interaction between bile acids and VLPs. In comparison to other bile acids tested, we observed the largest effect on VLP binding to cells using TLCA. Next, we pre-incubated VLPs with cholesterol with or without TLCA or GCDCA to investigate whether other bile components could enhance the effect of TLCA, as cholesterol is known to mediate entry and replication for many viruses, including GII.3 and GII.4 HuNoV(28, 34, 44, 45). We found that addition of cholesterol largely increased VLP binding, but only in the presence of bile acids (Fig. 3B), indicating a potential amplifying effect of bile acid-mediated binding in the presence of other hydrophobic molecules. Furthermore, removing cell membrane cholesterol by pre-treatment with the cholesterol sequestrant methyl-β-cyclodextrin (MβCD) reduced TLCA-mediated VLP binding in a dose-dependent manner, and at 2.5 mM completely abolished the TLCA-mediated binding (Fig 3C). Taken together, these results show that bile acids directly enhance VLP binding to cells, and that the presence of other hydrophobic molecules like cholesterol further enhance this effect.

**Figure 3.**
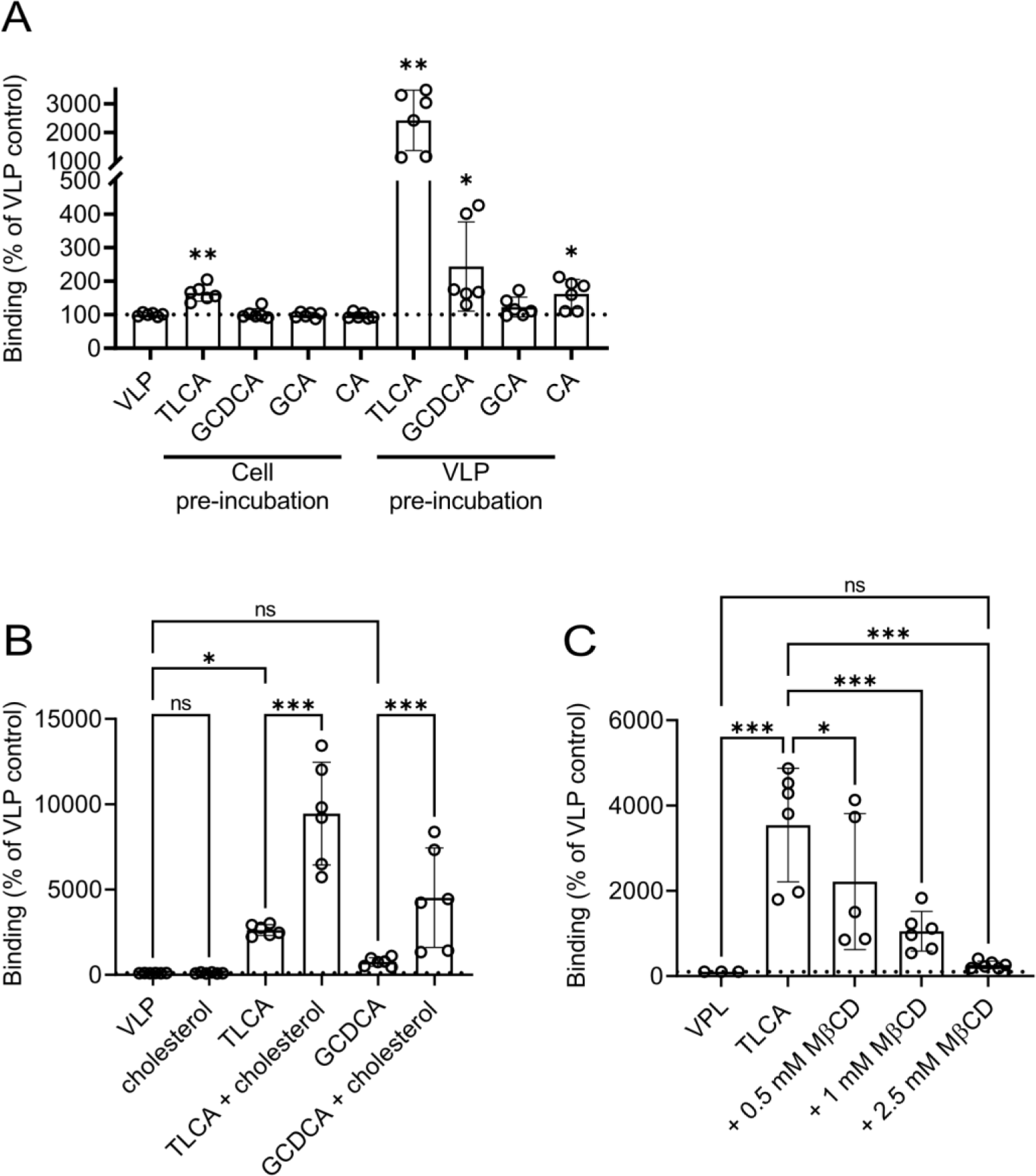
Flow cytometry analysis of TLCA-mediated HuNoV GII.4 VLPs binding to HuTu-80 cells in the presence of cholesterol. (A) VLP binding to HuTu-80 cells in the presence of bile acids where bile acids were either pre-incubated with cells before addition of VLPs (Cell pre-incubation), or pre-incubated with VLPs before addition to cells (VLP pre-incubation). (B) VLP binding to HuTu-80 cells after preincubation with TLCA, GCDCA, cholesterol or a combination of cholesterol and TLCA/GCDCA, respectively. (C) VLP binding to HuTu-80 cells in the presence of TLCA, after cell pre-treatment with cholesterol sequestrant MβCD. Binding was measured as median fluorescence and presented as percent of control (VLP binding in absence of bile acids). Data are from three independent experiments, represented as means ± SD. Statistical significance was determined with two-way ANOVA; *, P < 0.05; ***, P < 0.001.

### Bile acids are required for HuNoV VLP entry in HuTu-80 cells

As bile acids increased binding of HuNoV VLPs to HuTu-80 cells, we next investigated whether this also leads to uptake of VLPs into these cells. It has been shown that GCDCA indirectly increases entry of HuNoV GII.3 VLPs in human intestinal enteroids (34) through cellular effects, such as changes in endosomal acidification and apical membrane ceramide levels. However, it is not known whether human bile samples, or other bile acids than GCDCA, enhance entry, or whether bile acids can also mediate a direct effect on viral entry and uptake. We investigated this by looking at both binding and uptake in the presence of bile acids using confocal fluorescence microscopy. Alexa Fluor (AF)-488 labelled VLPs pre-incubated with bile acids were incubated with HuTu-80 cells on ice, followed by a one-hour incubation at 37°C to allow uptake. Live, non-permeabilized cells were stained with an anti-AF488 (green) primary and an AF568 (red)-conjugated secondary antibody to allow distinction between bound VLPs on the outside (red and green), and internalized VLPs inside (green only) the cells (as shown in Fig. 4A-D). We found almost no binding or uptake of VLPs in the absence of bile acids, as shown in Fig. 4A by low AF488 signal. However, when incubated with human bile (Fig. 4B) or hydrophobic bile acids (Fig. 4C-D), HuNoV VLPs both bound to (increased external green signal) and entered (increased internal green signal) HuTu-80 cells. We ascertained VLP uptake by careful analysis of single cells at variable z-stack positions. We found both single stained (green) internalized particles and double-stained bound particles at the cell membrane (orange) (Fig. 4E). We used these images to perform a relative comparison of total VLP signal in absence or presence of bile or bile acids. In the absence of bile and bile acids a very low VLP signal was detected (Fig. 4F). We next quantified VLP uptake for each condition and found that bile, GCDCA and in particular TLCA significantly increase VLP uptake (Fig. 4G), in line with signals seen in the images (Fig. 4B-D). Taken together, this suggests that bile acids are able to mediate direct binding to and subsequent uptake of HuNoV VLPs into HuTu-80 cells.

**Figure 4.**
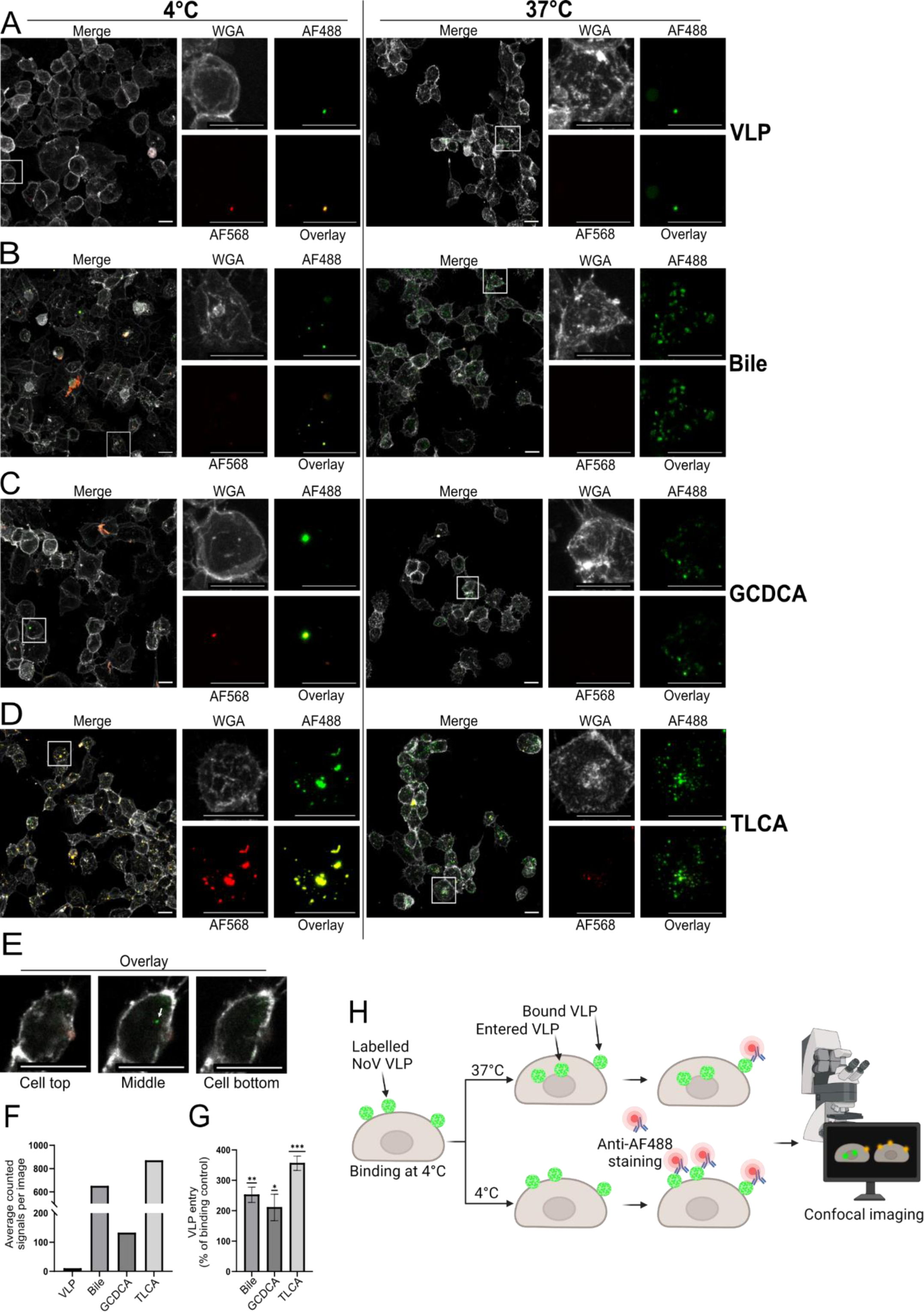
Confocal imaging showing bile acid dependent uptake of HuNoV GII.4 VLP into HuTu-80 cells. VLPs labelled with Alexa Fluor 488 were pre-incubated with bile or bile acids before addition to cells, after which they were allowed to bind (4°C) and subsequently enter cells (37°C). Remaining cell bound particles were stained with an anti-Alexa Fluor 488 antibody and a secondary Alexa Fluor 568 antibody. Cell membrane were stained with wheat germ agglutinin. Cells were visualized using a Leica SP8 confocal microscope and depicted are (A) representative maximum intensity projections of only VLPs, (B) VLP + human bile sample 3, (C) VLP + GCDCA and (D) VLP + TLCA. Scale bar for all images is 15 µm. (E) representative z-stack slices of one cell at increasing z-stack depth, visualizing VLP signal appearing inside of the cell (white arrow), never outside of the cell membrane. (F) Virus binding was increased in presence of bile or bile acid, assessed by object count of Alexa Fluor 488 signal which was measured for all images and displayed as the average number of objects per image. As the average number of objects per image for VLP only samples were very low, these were excluded from uptake quantification. (G) Virus uptake was quantified by measuring all virus particles (AF488) and extracellular virus particles (AF568). Increase in entry was measured as a relative comparison between extracellular and all virus object count for each sample. Uptake is shown as % increase in 37°C samples compared to the corresponding binding controls. (H) Graphical description of the method. Alexa fluor-488 labelled VLPs were allowed to bind to cells on ice before incubation at 37°C to allow for uptake (or on ice as control). Staining with anti-Alexa fluor 488 on non-permeabilized cells allows for separation between entered and non-entered VLPs. Statistical significance was determined with Welch’s *t* test; *, P < 0.05; **, P < 0.002; ***, P < 0.001.

### Bile acids and bile aggregate norovirus like particles

During confocal image analysis, we observed a bile acid-dependent increase of overall virus signal as well as the appearance of larger signal accumulations. The reason for these accumulations could not be determined from confocal microscopy, but indicated aggregation of VLPs, possibly resulting from interactions with the hydrophobic bile acids. We used negative stain electron microscopy to visualize VLPs mixed with bile acids to investigate this hypothesis. Control VLPs without bile or bile acids showed an even distribution of particles on the grid with only small clusters with few VLPs (Fig. 5A). The addition of bile acid GCDCA did not visually change the distribution, and no large clusters were observed either at pre-incubation concentration (4 mM) or binding concentration (250 µM) (Fig. 5B and C). However, in the presence of TLCA or human bile, we observed a marked difference in particle distribution. At 4 mM (pre-incubation condition), TLCA caused formation of large clusters consisting of several dozen VLPs (Fig. 5D), and even at the lower binding concentration of 250 µM, TLCA induced considerable accumulations with ten or more of VLPs (Fig. 5E). We used bile from patient number 3 as this sample yielded the largest increase in VLP binding to HuTu-80 cells (Fig. 1C). In the presence of 12.5% bile (pre-incubation concentration) and 0.8% bile (binding concentration) multiple VLPs clustered to a size similar to the TLCA conditions (Fig. 5F and G) and very few VLPs were seen as single particles. VLP clusters after bile treatment were more similar to the cluster size seen for TLCA than for GCDCA. As TLCA is more hydrophobic than GCDCA, this particular bile sample could contain more hydrophobic bile acids or components. These results suggest that TLCA, a highly hydrophobic bile acid, increases cellular attachment and VLP virus uptake by aggregation of HuNoV VLPs.

**Figure 5.**
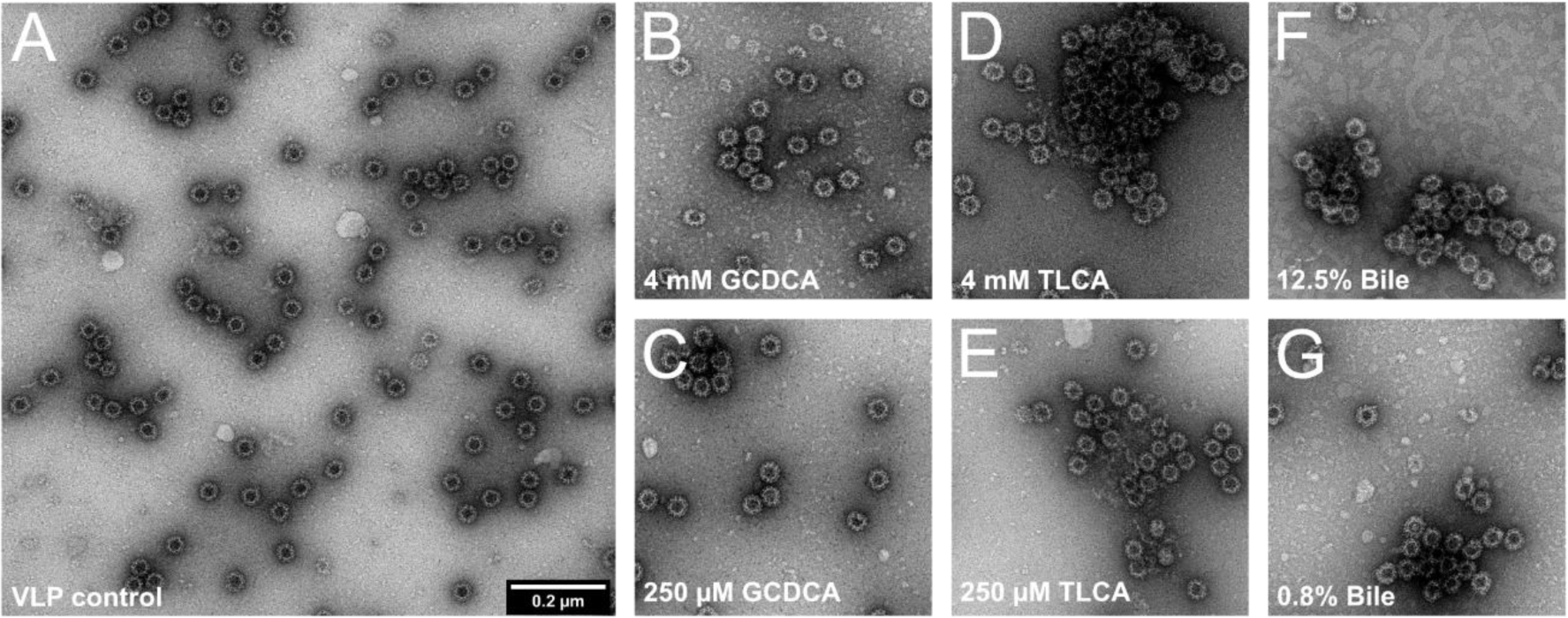
Negative stain of HuNoV GII.4 VLPs in the presence of bile acids and bile. HuNoV GII.4 VLPs imaged with electron microscopy, either (A) without the addition of bile or bile acids; (B) in the presence of 4mM or (C) 250µM GCDCA; (D) in the presence of 4mM or (E) 250µM TLCA; or in the presence of (F) 0.8% or (G) 12.5% human bile. Pictures shown are representative areas of each sample.

## Discussion

We demonstrate that bile acids mediate direct HuNoV GII.4 VLP binding to and uptake into a human duodenal cell line. This complements the knowledge that bile acids exert indirect effects to overcome the entry restriction of human norovirus replication in human enteroids. Additionally, we observe an accumulation of viral particles when incubated with human bile and hydrophobic bile acids, which has not been described before. When visualizing VLPs through EM (Fig. 5), the presence of bile and specific bile acids clearly results in viral particle accumulation, in particular with the highly hydrophobic TLCA (Fig. 5C and F). As some of these accumulations appear large, we propose that this could affect VLP cell entry, as our results suggest that accumulations do not inhibit viral uptake but rather increase it. It has been shown that enteric viruses such as rotavirus and human norovirus are secreted in extracellular vesicles, and that these vesicle-cloaked viral clusters enhance both infection efficiency and disease severity (40). Cell entry through vesicles mean that several virus particles can infect a single cell at the same time, which allows receptor-independent routes and increases the probability of successful replication (40, 41, 46–49). Still, in some cases of vesicle-mediated entry, infectivity is prevented by receptor-blocking; murine norovirus 1, hepatitis A virus and poliovirus released within extracellular vesicles are partially or almost completely blocked by antibodies against their respective receptors CD300lf, TIM-1, and CD155 respectively (40, 41, 48). However, as seen for vesicle-contained JC polyomavirus, the infectivity seems to be receptor-independent (49), explaining why the virus is able to invade tissues *in vivo*, which are otherwise less susceptible to free virus. Still, vesicles do not appear to completely overcome the requirement of receptor engagement, suggesting the possibility that they can be fully or partially released from vesicles in proximity to the cell surface, or possibly that they express markers that target them to receptor-expressing tissues, as speculated by Santiana *et al*. about murine norovirus, and Chen *et al*. regarding poliovirus (40, 41). Bile acids form micelles in the human intestine which aid in both excretion and absorption (36). Our results suggest a mechanism whereby TLCA and other hydrophobic bile acids, possibly in conjunction with cholesterol, provide a platform for aggregation of viral particles. We propose that aggregates may enter host cells by similar mechanisms as single particles, or by novel, hitherto unknown mechanisms.

Additionally, we show that hydrophobic bile acids are the most efficient enhancers of viral binding and uptake and that other hydrophobic molecules such as cholesterol can add to this effect (Fig. 3B) through an indirect mechanism not yet characterized. These results are consistent with previous studies that have shown effects of cholesterol on GII.3 as well as GII.4 replication (28, 34). The average human bile pool only contains trace amounts of highly hydrophobic bile acids like TLCA due to their cytotoxicity at higher concentrations (36, 50). However, human bile does not only contain bile acids, but also amphiphilic phospholipids and cholesterol (among other components) that together form mixed micelles in the duodenum (51). The strong, additive effect of cholesterol together with TLCA-mediated binding indicates that even a small amount of TLCA within a micelle could be enough to facilitate binding and uptake of HuNoV. We further showed that removal of cholesterol from the cell membrane with the sequestrant MβCD completely abolished the effect of TLCA, suggesting that cell membrane-associated cholesterol can enhance bile acid-mediated interactions. It can, however, not mediate VLP binding on its own, which parallels previous work by Murakami *et al.* showing that detergents (with similar properties as bile acids) do not mediate increased HuNoV replication (34). Thus, the increased binding and uptake cannot be explained by hydrophobic interaction alone and shows that bile acids seem to have unique features that make them pro-viral for HuNoV. Cholesterol is an important component in lipid rafts on the cell membrane, acting as platforms for receptor clustering, exploited by some viruses to enter cells. Several such virus-receptor interactions are abolished when disrupting lipid raft formation, such as Vaccinia virus interaction with CD98 (52), and Human herpesvirus 6 interaction with CD46 (53). In a recent study, GII.4 replication in FUT2 expressing human intestinal enteroids was inhibited by MβCD treatment, despite the fact that no bile acids were present (28), suggesting that cholesterol might be important for multiple receptor interaction mechanisms. It is therefore tempting to speculate that receptor clustering by cholesterol mediated lipid raft formation might also be true for HuNoV interactions virus host interaction.

In conclusion, we show that bile acids mediate direct attachment and entry of HuNoV GII.4 VLPs. We suggest that this mechanism is associated with the ability of bile to accumulate VLPs, which could be of relevance as infection of other intestinal viruses benefit from cell entry in clusters. Our results also indicate that other tissue fluid compounds such as cholesterol enhance the effect of bile acids on norovirus attachment and cell entry. Studying the properties of micelles isolated from norovirus-susceptible individuals might elucidate whether single bile acids like TLCA, or the overall bile acid composition is important for the increased binding and uptake. Taken together our findings provide valuable knowledge regarding norovirus biology and add to the understanding of attachment and entry of one of the most common causes of viral gastroenteritis.

## Acknowledgements

This research was supported by a research grant (2019–01472) from the Swedish Research Council, the medical faculty strategic research grant at Umea University (Dnr. FS 2.1.6-762-18), the medical faculty funded time for research at Umea University (Dnr. FS 2.1.6-2198-20) and Region Vasterbotten (ALF, Agreement for Medical Education and Research, Dnr HSN 86-2020).

We acknowledge the Biochemical Imaging Center at Umeå University and the National Microscopy Infrastructure, NMI (VR-RFI 2019-00217) for providing assistance in microscopy. The authors declare that they have no competing financial interest.

## Author contributions

E.P. and K.D. performed the binding experiments. E.P. and M.B. performed uptake assay and the confocal microscopy. K.D did initial sample preparation and L.S. and H.T performed the election microscopy. N.A. and A.L. proposed the project, designed the study and analyzed and interpreted the experiments. E.P., K.D., N.A., A.L. and M.B. wrote the manuscript. S.N. produced the HuNoV GII.4 VLPs. D.Ö. provided donated patient samples of human body fluids.

